# Including Crystallographic Symmetry in Quantum-based Refinement: Q|R#2

**DOI:** 10.1101/827170

**Authors:** Min Zheng, Malgorzata Biczysko, Yanting Xu, Nigel W. Moriarty, Holger Kruse, Alexandre Urzhumtsev, Mark P. Waller, Pavel V. Afonine

## Abstract

Three-dimensional structure models refined using low-resolution data from crystallographic or electron cryo-microscopy experiments can benefit from high quality restraints derived from quantum chemical methods. However, non-periodic atom-centered quantum chemistry codes do not inherently account for nearest neighbor interactions of crystallographic symmetry related copies in a satisfactory way. Herein, we have included these nearest neighbor effects in our model by expanding to a super-cell, and then truncating the super-cell to only include residues from neighboring cells that are interacting with the asymmetric unit. In this way our fragmentation approach can adequately and efficiently include the nearest neighbor effects. We have shown previously that a moderately sized X-ray structure can be treated with quantum methods if a fragmentation approach was applied. In this study, we partition a target protein (4gif) into a number of large fragments. The use of large fragments (typically hundreds of atoms) is tractable when a GPU based package such as TeraChem is employed or cheaper (semi-empirical) methods are used. We run the QM calculations at the HF-D3/6-31G level. We compare and contrast the models refined using a recently developed semi-empirical method (GFN2-xTB). To validate the refinement procedure for a non-P1 structure, we use a standard set of crystallographic metrics. We show the robustness of our implementation by refining 13 additional protein models across multiple space-groups and present the summary of the refinement metrics.

**Synopsis:** C-terminal coiled-coil domain of transient receptor potential channel TRPP3 in the P321 space group (PDB code: 4gif) is re-refined with restraints from quantum chemistry using Hartree-Fock theory.

## 1. Introduction

In experimental structural biology, the atomic model (three-dimensional structure) of a bio-macromolecule is iteratively improved by a procedure known as refinement. In principle, refinement is a restrained or constrained optimization problem with respect to model parameters

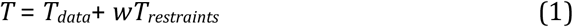

The target function (1) is a weighted sum of two components. T_*data*_ is derived from experimental data, *w* is an empirical scale factor, and T_*restraints*_ is *a priori* knowledge about the problem hereafter referred to as restraints. Typically, bio-macromolecules have many atoms and therefore the parameter space is high dimensional. This means that refinement requires a large number of steps to converge. Current refinement procedures (e.g. L-BFGS, Liu & Nocedal, 1989) need the gradient of (1) at each and every step of the minimization; therefore the computational cost can be intractable, especially for large molecules. To overcome this, the model parameterization in standard refinement is very simplistic and relies on parameterized libraries for chemical information that are used as restraints in (1) (for example, Afonine *et al*., 2012). Unfortunately, the traditionally employed parameterized restraints alone are not sufficient to provide information for accurate refinement using only low-resolution experimental data (Zheng *et al*., 2017a,b). Therefore, quality and reliability improvement of refined protein X-ray crystallographic structures is an on-going challenge, while developing methods that can provide better and more efficient restraints in a computational tractable fashion is an active area of research.

The idea of quantum refinement is based on the premise that the atomic model of a bio-macromolecular structure can be improved by employing restraints derived from quantum chemistry calculations. Quantum chemical *ab initio* restraints indeed have the ability to provide more accurate bio-macromolecular structures but at the cost of requiring far more computational resources. A way to overcome this difficulty is to use fragmentation methods that can effectively reduce large and complex systems (Zheng *et al*., 2017b) to smaller and more tractable units.

A number of different quantum refinement approaches are available. Ryde and co-workers (Ryde 2003; Ryde *et al.*, 2003; 2003; Nilsson *et al.*, 2004;) have focused their efforts on hybrid QM/MM based approaches (Senn & Thiel 2009), which concentrates the computational resources around a given site of interest. Merz and co-workers (Yu *et al.*, 2005; Yu *et al.*, 2006) have applied standard semi-empirical methods, presumably due to the inherent low resource scaling of their computational cost. Alternatively, we have initiated a fully *ab initio* quantum refinement using a fragmentation approach (Zheng *et al.*, 2017a,b). Each group has developed their own implementations and made them available as programs ComQum, DivCon, and Q|R, respectively. However, to date none of these quantum-based solutions have become standard in protein crystallography.

Our Q|R roadmap contains a list of challenges that need to be solved to provide a real alternative to traditional refinement. In this paper, we wish to address the next major issue — accurately treating nearest neighbors arising from the crystal symmetry. The PDB contains entries that were solved in many different space groups with P1 symmetry being far from the most common (figure 1). Therefore, we need a general and robust approach to properly account for crystallographic symmetry that works together with our fragmentation based-approach. Herein, we present our solution to this problem. Firstly, in order to account for the covalent and non-covalent interactions between symmetry related neighbors we expand the contents of the unit cell to construct a super cell. In a second step, we truncate the model based on the cut-off distance to the central unit cell to define a region that we term the ‘super-sphere’. The latter is then subdivided into the manageable fragments using interaction-based graphs, see Zheng *et al.* (2017b). We finally validate our method by refining a set of non-P1 structures and compare it with corresponding results of classic refinements.

**Figure 1.**
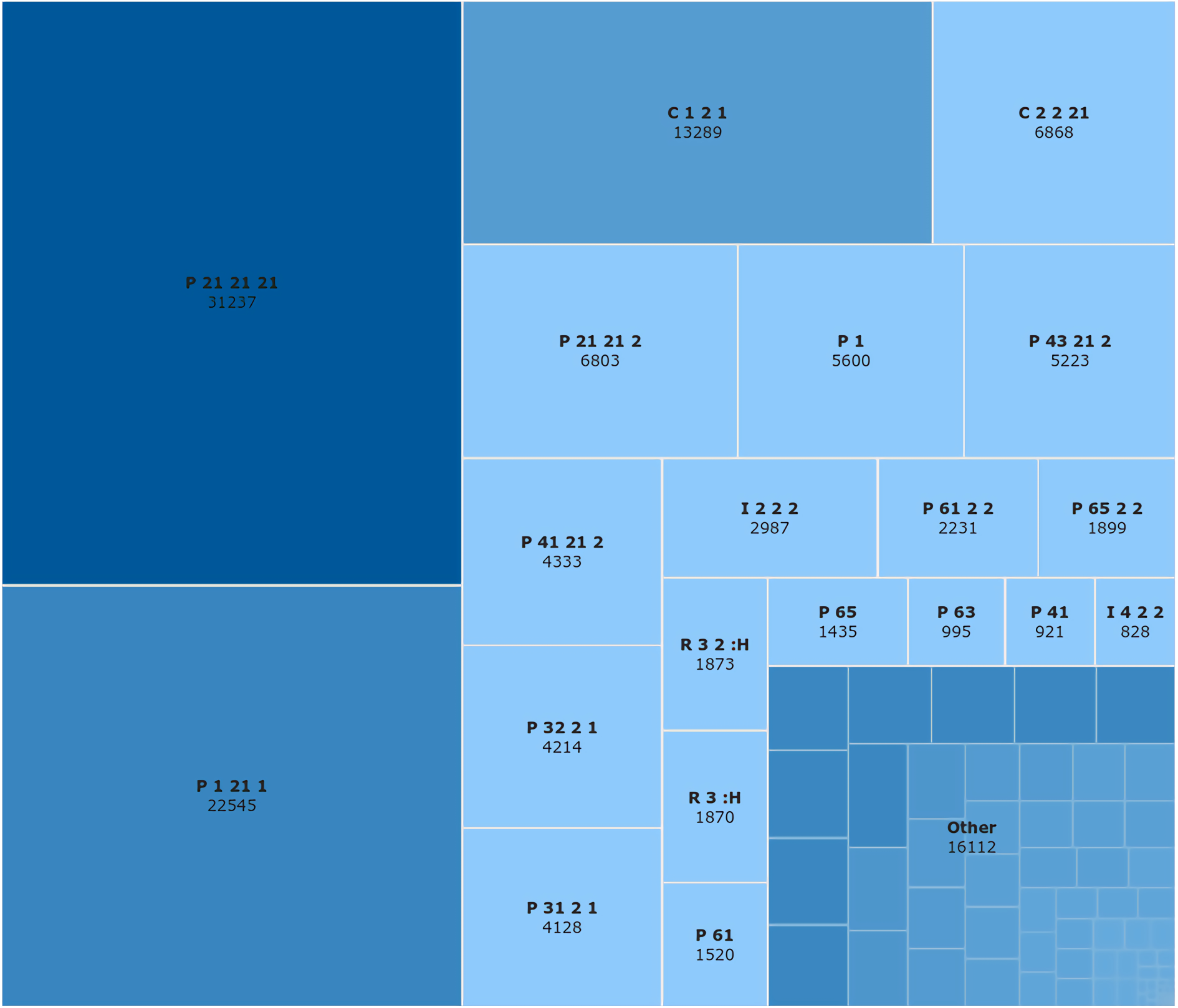
The distribution of space groups present in the entire PDB as of June 2019, where the number indicates the number of instances for each space group.

For simplicity, in what follows we refer to the QM calculations at the HF-D3/6-31G level using TeraChem software as *TeraChem (method, refinement)*. Likewise, we refer to calculations using semi-empirical GFN2-xTB method, as *XTB (method, refinement)*. Finally, calculations using standard parameterized libraries for stereochemical restraints as implemented in CCTBX package are referred to as *classic (standard)* or *CCTBX refinement (method)*.

## 2. Methods

### 2.1. Model selection and preparation

The choice of test examples used to validate our new method accounting for crystallographic symmetry in quantum-based refinement is rather simple; the symmetry should not be P1. Here, we selected an atomic model of C-terminal coiled-coil domain of transient receptor potential channel TRPP3 (PDB code: 4gif) that was reported to have P321 symmetry. The structure has a number of non-covalent interactions between symmetry copies that makes it attractive for this work. Also, the diffraction data resolution of 2.8 Å and rather poor data completeness at both low- and high-resolution ends makes the use of accurate restraints ever important. To further demonstrate robustness of our approach and the refinement procedure, 13 additional structures have been selected, covering the resolution range of 1.5-3.8 Å with symmetries P2_1_2_1_2_1_, P6_5_, P2_1_2_1_2, P4_3_2_1_2, P312, P6_4_, P6_4_22, P3_1_21, I432, P6_3_, P6_5_, I4_1_22 (PDB codes: 6nfs, 6n1L, 5zkt, 6ney, 6njg, 6agy, 6jqs, 5xsL, 3wap, 3hzq, 2r30, 3nr7, 5kzb).

An atom-complete and correctly protonated atomic model is required for quantum chemical calculations. Unfortunately, most crystallographic models in the PDB are atom-incomplete (for example, lack hydrogen atoms, side chains or parts thereof). Therefore each input model must be pre-processed before quantum refinement. For this, we have developed the *qr.finalise* tool that is part of Q|R software suite.

### 2.2. Handling neighbors: the crystal symmetry

In order to account for contributions to the target function (1) that arise from symmetry related copies, we have developed a computationally efficient algorithm. Provided the asymmetric copy of the molecule, the symmetry type and the distance cutoff, the procedure will first expand the asymmetric copy by applying all symmetry operators, and then truncate all atoms that fall outside the distance cutoff from the nearest atom in the asymmetric copy. Since QM calculations require atom-complete models, the truncation is done such that if an atom falls within the distance cutoff then the whole residue that contains this atom is preserved as the neighbor. The details of the algorithm are provided in the Appendix.

### 2.3. Quantum Refinement

We have been developing a software package called Q|R to perform quantum-based crystallographic refinement of bio-macromolecules (Zheng *et al*. 2017a, 2017b). Q|R interfaces to the CCTBX open source project (Grosse-Kunstleve *et al.*, 2002) for the computation of all additional quantities needed for the refinement such as *T*_*data*_ in (1). Also, Q|R interfaces to the ASE package (Bahn & Jacobsen, 2002) which contains wrappers to many modern QM computational packages that can be used to calculate the target and gradients needed as restraints in (1). To perform QM calculations in this work we use the semi-empirical method as implemented in XTB package (Grimme *et al*., 2017; Bannwarth *et al.*, 2019) and the Hartree-Fock (HF) method from TeraChem package (Ufimtsev & Martínez, 2009: Titov *et al*., 2013).

XTB is a software that uses an extended tight-binding semi-empirical method focused on the accurate prediction of reliable geometries, frequencies and non-covalent interactions (GFN-xTB) recently developed by Grimme (Grimme *et al*., 2017; Bannwarth *et al.*, 2019). We created a new ASE interface to Grimme’s XTB code^1^. We employ the second generation method denoted GFN2 that is currently the only semi-empirical method able to treat hydrogen bonds natively without an empirical correction due to the use of multipole-based electrostatics (Bannwarth *et al*., 2019). To improve the SCC (Self Consistent-Charge) convergence for the charged fragments a non-default electronic temperature of 500 K together with the implemented generalized Born solvation model (water parameters) is employed. Charged proteins, with zwitterionic chains, are inherently problematic for molecular orbital methods as the band gap can become very small which leads to problems finding a variational energy minimum. While sometimes a small dielectric constant is chosen to mimic the charge-screening effects from the rest-protein environment, we opted for a high dielectric constant (~80 for water) as a safe choice to ensure SCC convergence within our automated calculations.

TeraChem is a quantum chemical package (Ufimtsev & Martínez, 2009: Titov *et al*., 2013) that uses the graphical processing unit (GPU) as the computational device. The Hartree-Fock (HF) method with Grimme’s dispersion correction D3 (Grimme et al., 2010) was employed, based on recommendations from a benchmarking study from Goerigk *et al*. 2013. However, the fragmentation based-approaches employed in the present work allow for favorable scaling of QM computations without severe restrictions on the basis set size. Therefore, we opted for the dispersion-corrected HF-D3 method in conjunction with the 6-31G basis set (Hehre *et al*, 1972). The computations were performed accounting for polar (water) environment by means of the COSMO polarizable continuum model (Liu *et al*, 2015; Barone & Cossi, 1998; Truong & Stefanovich, 1995). In what follows we will refer to this method as HF-D3 or simply TeraChem.

The protein model needs to be divided into smaller fragments that can then be efficiently treated by standard quantum mechanical methods. The approach implemented in Q|R is detailed in Zheng *et al.* (2017b). Briefly, the whole protein is divided into disjoint parts called *clusters* using a *divide and conquer* based approach. In places where covalent bonds were cut (e.g. peptide bonds) we use hydrogen capping atoms to terminate the dangling bonds to satisfy valence requirements (Senn & Thiel 2009). Then we search for residues that are interacting with each fragment; the set of interacting residues for a given cluster is referred to as a *buffer* region. The buffer regions are important to reduce errors in the gradients for atoms near the edge of the cluster (boundary). The larger the buffer, the more accurate the gradients will be, but the amount of computational resources will also increase so it becomes a trade-off. A cluster combined with its associated buffer region is known as a *fragment*. A single gradient evaluation of the quantum chemical energy is made for each and every fragment. The quantum mechanically computed gradients for each cluster are then extracted from the fragments and added together to create a total gradient for the entire molecular system, while the gradients for the buffer region and capping atoms are cast away. To ensure this does not incur computational errors, we have developed an automated procedure to analyze the quality of fragment-derived gradients by comparing them with a reference gradient that is either calculated from the whole model or from a set of the largest possible fragments.

## 3. Results

All refinements were performed using Q|R v1.0. The fragmentation procedures for the 4gif model resulted in eight disjoint clusters. The XTB based refinements were sufficiently fast to enable us to trial several settings (such as number of minimization iterations, LBFGS optimization step length, maximum allowed number of residues per cluster) to explore the procedure in terms of runtime and accuracy. For the 4gif model a maximum of 15 residues per cluster was found optimal. The root-mean-square deviation of the gradient using the standard buffer (single-buffer) region is 0.6 kcal/mol/Å compared to a converged gradient using a notably enlarged buffer (triple-buffer). Finally, to provide an *ab initio* based refined structure of 4gif, a refinement using HF-D3 in TeraChem was performed.

Similar to our previous work (Zheng *et al*., 2017b), we used a generally accepted set of crystallographic and model statistic metrics (table 1) to validate and compare refinement results. We find overall *R* factors are very similar across all refinement results, which is not unexpected as we re-refined an already finalized published model. The lowest *R*_*free*_ result is from TeraChem. Furthermore, both QM based refinements produced a systematically smaller *R*_*free*_-*R*_*work*_ gap thus indicating less data overfitting. The TeraChem refined model also has the most improved secondary structure geometry of the refined model as indicated by the highest Ramachandran Z-score^2^. XTB and CCTBX refinement results have more negative Z values but within the reasonable range. The clashscore (Chen at al., 2010) for both TeraChem and XTB based refinements has been decreased by almost an order of magnitude. This is also expected as QM methods account for short- and-long range interatomic interactions, while classic refinements use only non-bonded repulsion terms. The MolProbity (Chen *et al*., 2010) score suggests that the overall quality of QM refined models is similar to models solved at resolutions around 1 Å, while the initial and classically refined models are more similar to models refined at 1.9 and 1.6 Å resolutions, respectively.

**Table 1.** Statistics for 4gif shown for initial model as obtained from the databank (PDB), as well as re-refined model using standard restraints (CCTBX), restraints from semi-empirical (XTB) and *ab initio* HF-D3 (TeraChem) methods. For consistency, *phenix.model_vs_data* and *phenix.model_statistics* were used to obtain the reported values in all four cases. All-atom root-mean-squared deviations (RMSD) were calculated between the initial model from PDB and three re-refined models using *phenix.superpose_pdbs*. The same input model preconditioned for quantum refinement as described in §2.1 was used in all three refinements.

| Model |  | PDB | CCTBX | XTB | TeraChem |
| --- | --- | --- | --- | --- | --- |
| Metric |  |  |  |  |  |
| R-factor | work | 0.2335 | 0.2443 | 0.2574 | 0.2473 |
|  | free | 0.2994 | 0.2992 | 0.2941 | 0.2910 |
|  | gap | 0.0658 | 0.0549 | 0.0367 | 0.0437 |
| RMSD | Bonds (Å) | 0.014 | 0.007 | 0.011 | 0.015 |
|  | Angles (°) | 1.65 | 0.83 | 1.52 | 1.60 |
| Rama. plot (%) | Favored | 100 | 100 | 100 | 100 |
|  | Outliers | 0 | 0 | 0 | 0 |
|  | Z-score | -2.85 | -1.84 | -0.57 | 2.72 |
| Rotamer outliers (%) |  | 5.26 | 0 | 2.63 | 0 |
| Clashscore |  | 6.92 | 11.05 | 1.38 | 1.38 |
| C<sub>β</sub> deviations |  | 0 | 0 | 0 | 0 |
| MolProbity score |  | 1.93 | 1.61 | 1.19 | 0.87 |
| RMSD (Å) |  | n/a | 0.370 | 0.412 | 0.389 |

Since correct handling of crystal symmetry is the focus of this work, we annotated and analyzed all hydrogen bonds (H-bonds) on the interface between the molecule in the asymmetric unit and all surrounding symmetry copies – 14 bonds in total. Figure 2 shows the distribution of hydrogen-acceptor (H-A) distances and donor-hydrogen-acceptor (D-H-A) angles. Hydrogen bonds range in strength greatly (for example, Steiner, 2002). In particular, the values for distances are expected to vary between 1.2 and 3 Å, clustering around 1.6-2.0 Å; angles can be in 90-180 degrees range, with strongest bonds approaching the linear configuration. Refinement results using both XTB and TeraChem produce models with the most hydrogen bonds having their parameters in the expected range and with most values indicating stronger bonds. This not only shows the superiority of QM-based refinements compared to classic refinement, but also proves that our algorithm can handle crystal symmetry in the refinement procedure correctly. We also note that H-bond parameters for classic refinement cover a much broader range of sterically possible values with less inclination to favor a particular value or distribution. This is because classic refinement is agnostic to this kind of interactions and only includes non-bonded repulsion terms that are counter-productive for H-bonds.

**Figure 2.**
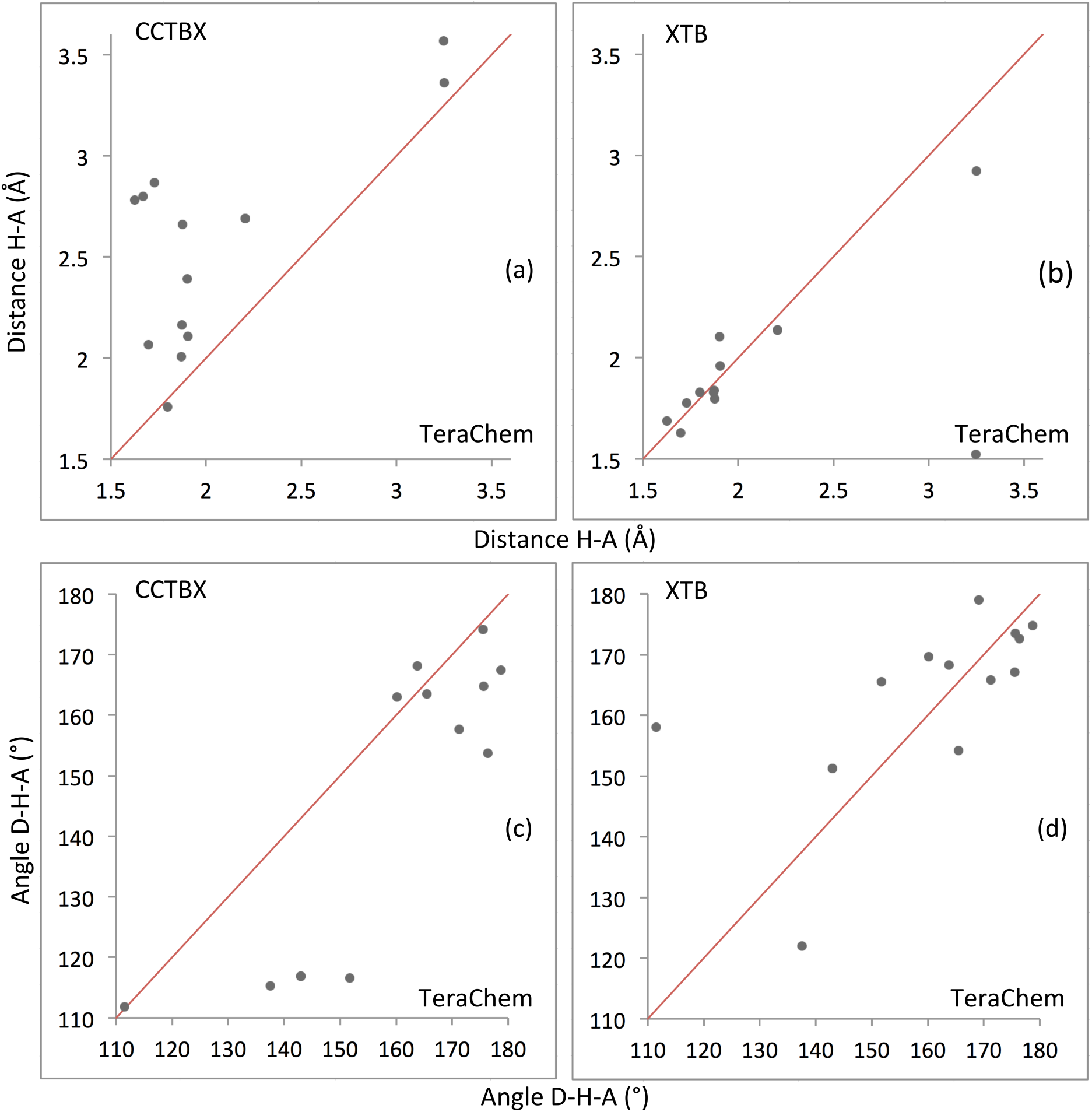
Distribution of hydrogen bond parameters for all 14 cross-symmetry bonds in 4gif: (a, b) hydrogen-acceptor (H-A) distances and (c, d) donor-hydrogen-acceptor (D-H-A) angles. Scatter plots contrast HF-D3 (TeraChem) refined models versus classic (CCTBX) and XTB results.

A remarkable example is the water S4 that bridges Glu718 with the main chain oxygen O of symmetry related Arg713 (figure 3). In both hydrogen bonds, the CCTBX method has the longest bond length with the TeraChem method having the shortest. The charge on the acid moiety changes the nature of the bond but it remains in the moderate strength category. Hydrogen atoms are effectively absent in the CCTBX refinement because when protonating the water-oxygens the occupancy of the hydrogens is set to zero and they only participate through non-bonded repulsion. The protonation algorithm places the hydrogen atoms in the same arbitrary orientation for all water molecules so any hydrogen bonding (or lack thereof) is based on the chemical restraints component of the target function.

**Figure 3.**
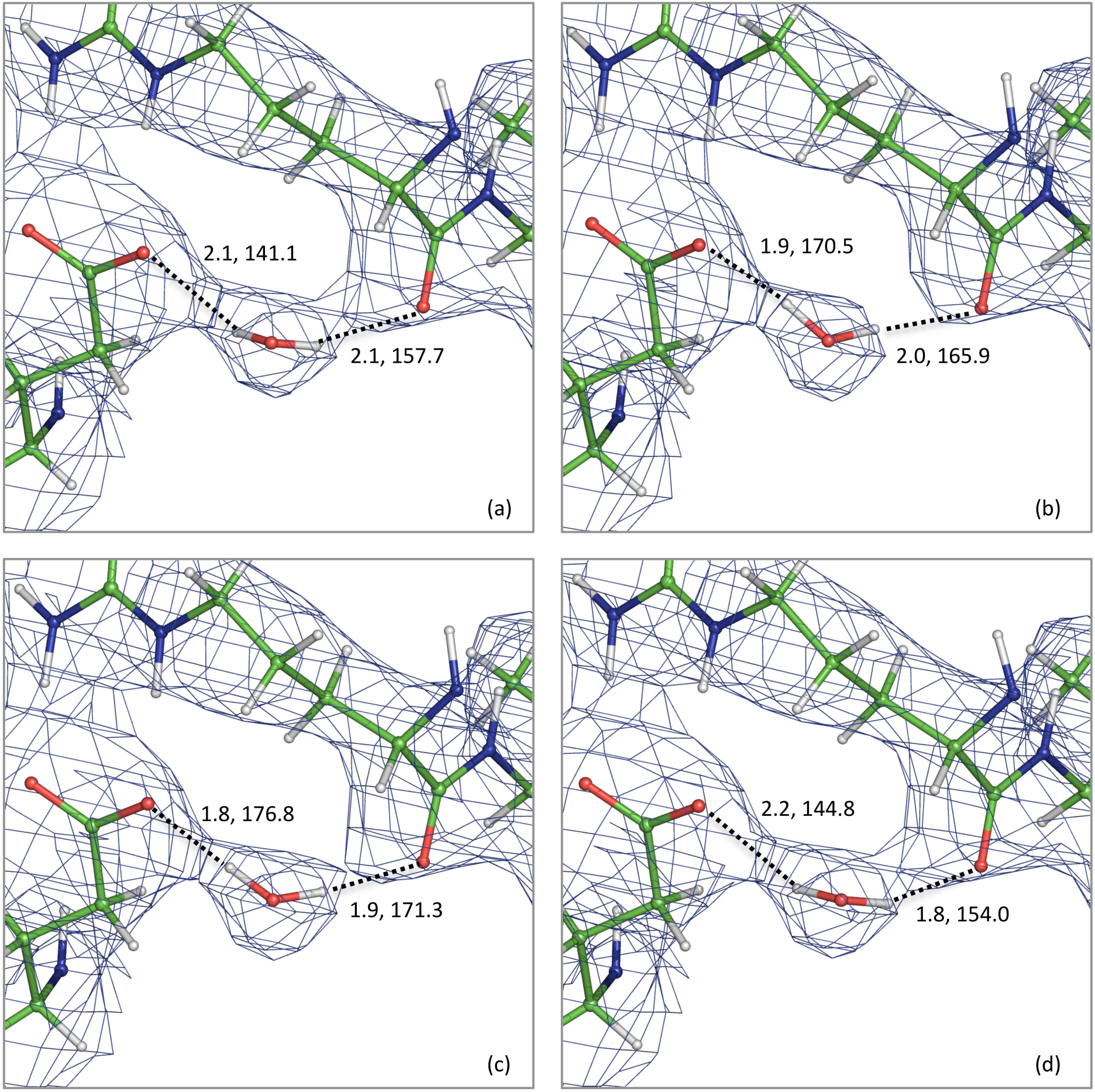
Water S4 is coordinated via both hydrogen atoms to oxygen atoms of the protein. One is hydrogen bonded to the carboxylic acid of a symmetry copy of Arg713. The other is coordinated to an acid side-chain oxygen atom (OE2) of Glu718. Pairs of numbers indicate bond distance in Ångström (dash line) and the angle in degrees based on water oxygen, corresponding water hydrogen and protein oxygen. Panels show model before refinement (a) and refinement results for (b) XTB, (c) TeraChem and (d) CCTBX.

To avoid basing our conclusions on a single test case and also to exercise and test our refinement protocol further, we performed quantum refinement with the XTB code for additional 13 structures selected from the PDB. They represent a rather broad range of resolutions and space groups (see § 2.1 for details). Overall, we observe the same trend in refinement statistics: R-factors remain the same or improve only slightly (figure 4) and model geometry improves the most for QM refinements (not shown). Figure 5 shows the distribution of H-bond parameters (same as for 4gif: bond length and angle) for all bonds across symmetry gathered in all 13 models. One can see that the XTB-refined models contain more (higher histogram) and stronger hydrogen bonds (shifted to lower distances). This is encouraging, as it shows the benefit of quantum methods that explicitly handle hydrogen-bonding interactions.

**Figure 4.**
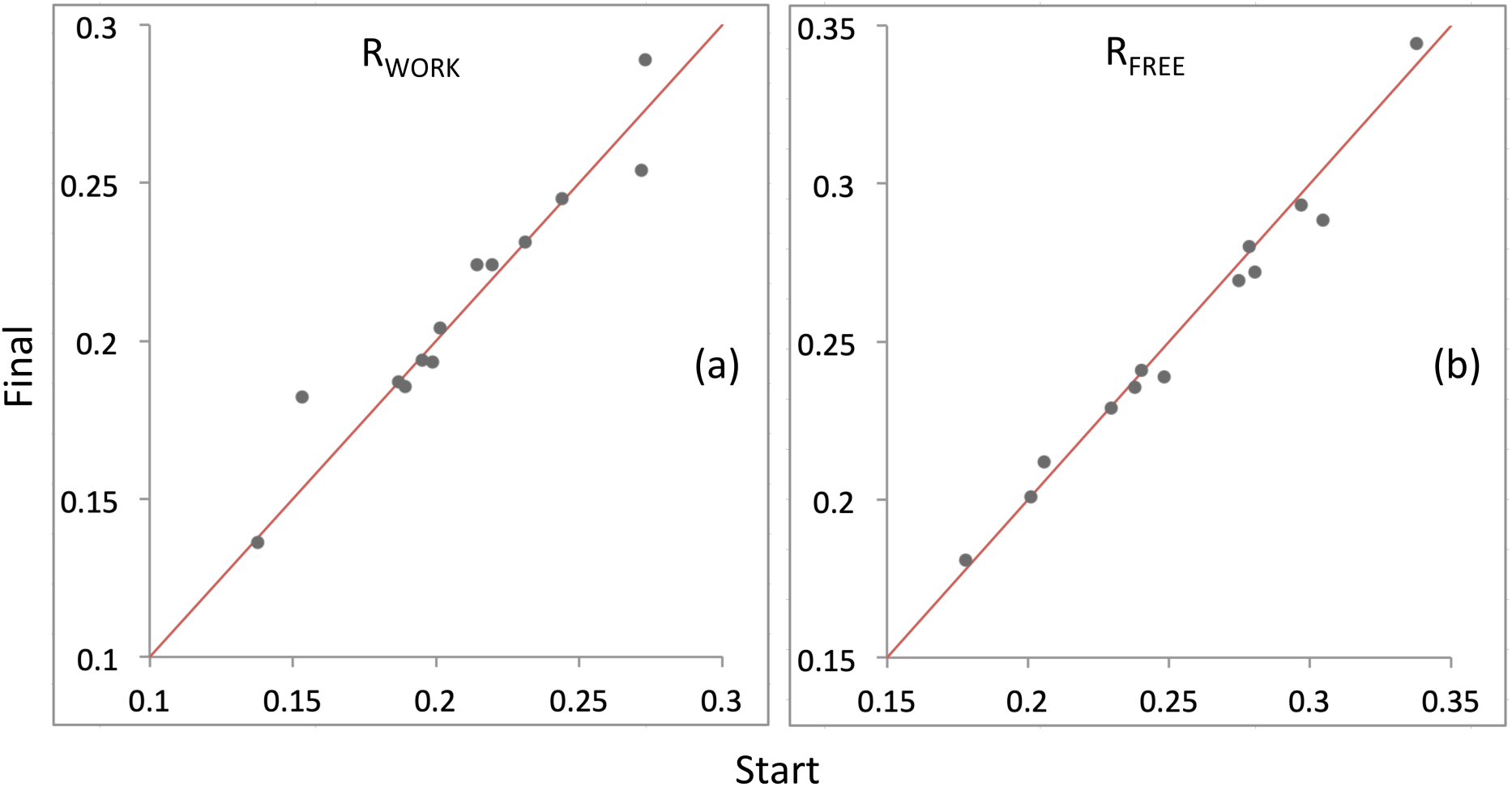
*R*_*work*_ and *R*_*free*_ summary for all 13 re-refined PDB models shown before and after XTB refinement.

**Figure 5.**
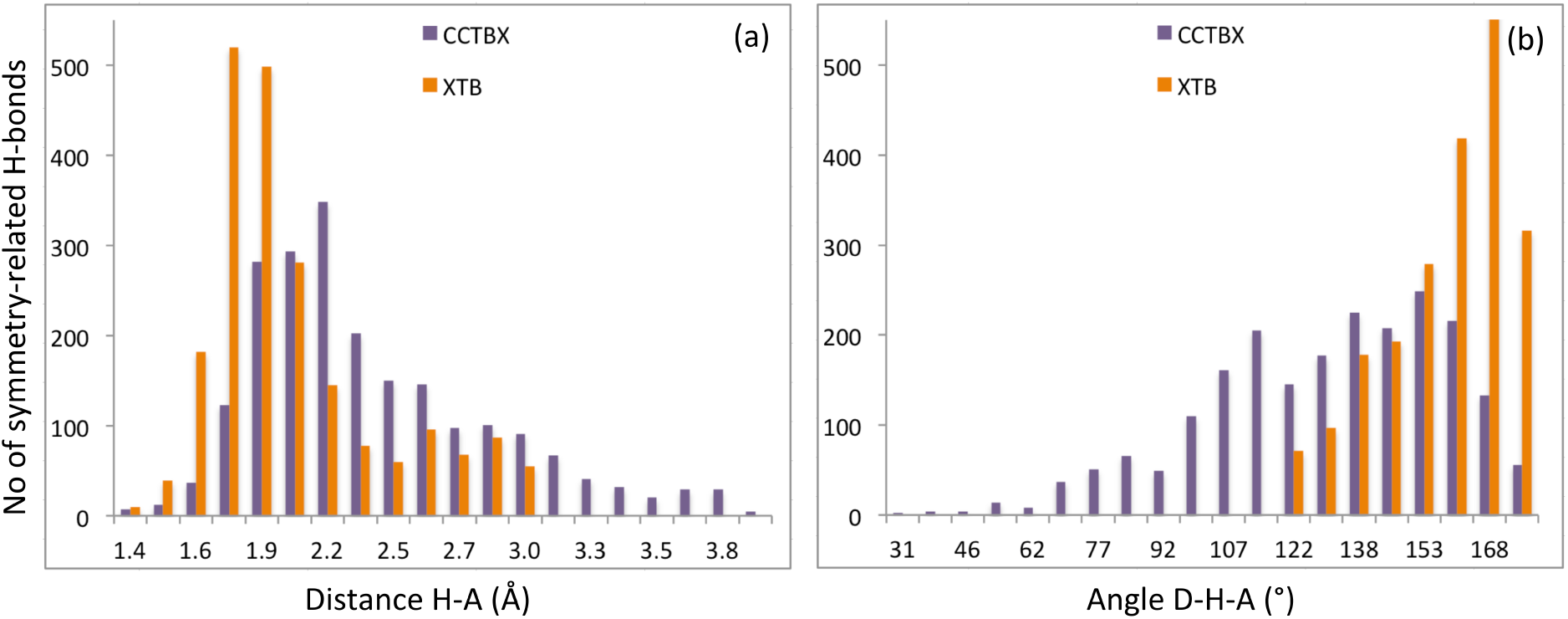
Distribution of hydrogen bond parameters across symmetry copies for all 13 re-refined PDB models: (a) hydrogen-acceptor (H-A) distances and (b) donor-hydrogen-acceptor (D-H-A) angles.

## 4. Conclusion

In this work we set out to design, implement and validate a method for including crystallographic symmetry into our quantum refinement procedure Q|R. We have achieved this by expanding the unit cell to include nearest neighbors in the quantum chemical calculations. In order to perform these calculations more efficiently, we use a truncated model called a super-sphere to remove all non-interacting residues from the symmetry copies in the fully expanded super cell.

To tune the performance of our implementation, we interfaced a new semi-empirical method known as GFN2-xTB as it offers an improved refinement over the standard CCTBX, but at a fraction of the cost of traditional QM methods such as Hartree-Fock. The Q|R codebase was designed to be as modular as possible, which makes it trivial to adapt to new quantum chemical programs to obtain energy functions and/or/only their gradients.

To the best of our knowledge, this is the first *ab initio* quantum refinement of a crystal structure to be carried out using quantum restraints that included nearest neighbor effects. We have re-refined the 4gif structure and presented key structural metrics to assess the quality of the quantum-based refinement. In addition, 13 other structures were also re-refined to demonstrate the robustness of our implementation. The next step on our road map is to handle alternate locations, and we will further explore the potential of quantum-based refinement for X-ray crystallography and cryo-EM in our on-going development of the Q|R project.

## Acknowledgments

This work was enabled by the financial support from the National Natural Science Foundation of China (Grant No. 31870738) and support from the Shanghai Eastern Scholar Program. AU acknowledges the support and the use of resources of the French Infrastructure for Integrated Structural Biology FRISBI ANR-10-INBS-05 and of Instruct-ERIC. HK acknowledges support by the ERDF project SYMBIT (CZ.02.1.01/0.0/0.0/15_003/0000477).

## Appendix

## 1. Preamble

Let (*X*_*m*_, *Y*_*m*_, *Z*_*m*_), *m* = 1, *M* be the Cartesian coordinates of *M* atoms of a model under study. This model, composed of *J* residues (or other atomic groups), is considered to be in some crystal, and {***r***_*m*_} = {*x*_*m*_, *y*_*m*_, *z*_*m*_}, *m* = 1, *M* are corresponding crystallographic (fractional) coordinates. We note that the model does not necessarily belong to the unit cell closest to the origin, that we call below *basic unit cell*, i.e. that the conditions

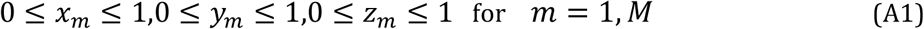

are not necessarily satisfied for all atoms. It is rather usual that a fraction of atoms stick out of this box and a model belongs to several unit cells as Figure A1 illustrates schematically.

There is a set of crystallographic symmetries *S*_*k*_ = {*R*_*k*_; ***T***_*k*_}, *k* = 1, *K* and eventually a set of non-crystallographic (local) symmetries, *S*_*l*_ = {*R*_*l*_; ***T***_*l*_}, *l* = 1, *L*, that are applicable to the given model. Here *R* stands for the rotation matrices and **T** for the translation vectors expressed in fractional coordinates. In the crystal, the model, that we will call the *original* copy, may interact (and usually does) with some of its symmetry-related copies. We define interaction as a presence of atoms at a distance to at least one of atoms of the original copy shorter than a given threshold value *d*_*contact*_. From all symmetry-related copies we want to select all residues that contain at least one of such interacting atoms. Also, we check atom overlaps, i.e. the situation when an atom after applying a symmetry operation is closer than *d*_*overlap*_ to an atom in the original copy. Default values are *d*_*contact*_ = 3 Å and *d*_*overlap*_ = 1 Å.

## 2. Model and data preparation

### 2.1. Model deorthogonalization

First, the atomic Cartesian coordinates, e.g. taken from PDB, are deorthogonalized (converted into fractional coordinates). The result of this operation is needed for two goals: 1) to generate symmetry related copies and 2) to compare the *original* model with these generated copies.

If the local (non-crystallographic) symmetry operations are defined, they also should be in form of rotation matrices and translation vectors to be applicable to the fractional coordinates of the given deorthogonalized model.

### 2.2. Symmetry preparation

We prepare the full list of *KL* symmetries that will be applied to each atom of the original copy:

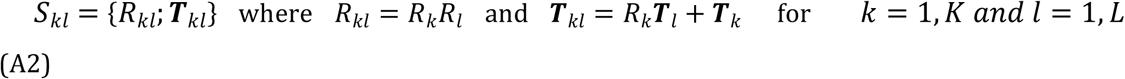

The local symmetry operations are applicable only to the given copy. For this reason we do not shift the original model to the basic unit cell even when this could simplify some further analysis.

If local symmetries are absent, the (A2) list simply consists of crystallographic symmetries.

### 2.3. Box annotation with original model

For comparison, it is convenient to have a copy in the basic unit cell. Since the original model may be extended over several unit cells, we identify all these cells. For short, we call them *boxes* and refer to them by their ‘corner’ that has the minimal coordinate values; by definition, all of them are integers. For example, the basic unit cell is the box (0,0,0). To identify all the boxes containing the original model we do the following steps.

a. We calculate minimal and maximal values for each of the fractional atomic coordinates,

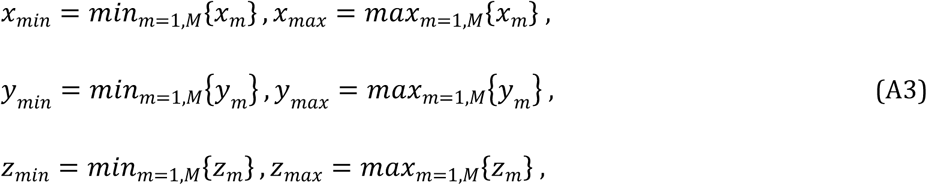
b. We define minimal and maximal *integer* translations **t** by each coordinate that may put some model atoms inside the basic unit cell,

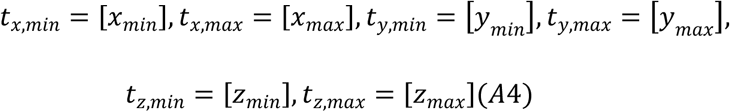

where [x] stands for the floor operation that defines the largest integer less than or equal to x. These parameters define a super-cell that covers the whole original model. This collection of boxes may or may not contain the basic cell, depending on where the model is happened to be in space. Parameters in (A4) define the origin and the farthermost corner of this super cell.
c. We rename the boxes that compose the super-cell in some sequential order, 1 ≤ *n* ≤ *N*. We note by ***t***_*n*_ = (*t*_*x,n*_, *t*_*y,n*_, *t*_*z,n*_) coordinates of the ‘left bottom’ corner of this box; they are integer numbers. There are in total

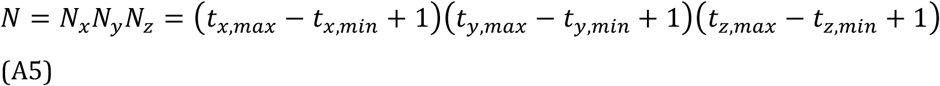

boxes even when some of them may actually contain no atoms (e.g. if the model is ‘diagonal’, corner boxes most distant from the ‘diagonal’ may be empty; Fig. 2). Subtracting (*t*_*x,n*_, *t*_*y,n*_, *t*_*z,n*_) from the coordinates of atoms that belong to the box *n* shifts these atoms into the basic unit cell with <u>*all*</u> coordinates belonging to [0,1] range. Applying different shifts to the content of different boxes means that the shifted model may appear as “torn apart” when inspecting on graphics. This is different from shifting the entire model to maximally belong to [0,1] box, in which case some atoms of such model can still protrude outside the [0,1] range (Figure A1).
d. To gain computing time, for further calculations a *box content table* composed of *M* lines and two columns is created. Initially the first column contains consecutive numbers and the second contains the number of the box this atom belongs to. Then this table is reordered by the box number, 1 ≤ *n* ≤ *N*, so that all lines for the same box are neighboring to each other.

Below, we will generate symmetry related copies from the full model but will compare them independently with the content of each of the *N* boxes defined above, one by one.

### 2.4 Model to generate symmetry related copies

The same *original* deorthognalized model, with no shifts yet, will be used for another goal, which is to generate its symmetry related copies. To make the procedure faster and more efficient, we prepare an intermediate object.

For each residue *j* we determine and save the fractional coordinates of its geometric center ***C***_*j*_ calculating the mean value by each coordinate of all atoms of the group. We also calculate the radius *r*_*j*_ of the minimal sphere centered on ***C***_*j*_ that covers all atoms of this residue. In other words, *r*_*j*_, in Å, is the maximal distance from ***C***_*j*_ to the atoms of this residue. Then, using the contact distance *d*_*contact*_ that is a parameter of the problem, we define the width of margin, also in Å,

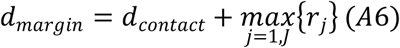

to be used to build the mask of the model for comparison.

## 3. Search for the neighboring symmetry related copies

### 3.1. Masks preparation

Search for the symmetry copies that make a contact with the original model is done as a cyclic procedure over *N* boxes defined above. First, for the box *n* and the group of atoms of the original model that belong to this box we prepare a mask as follows:

a. Define *extended basic unit cell* such that the fractional coordinates of its origin and farthermost corner are (-*d*_*margin*_/*a*, -*d*_*margin*_/*b*, -*d*_*margin*_/*c*) and (1+*d*_*margin*_/*a*, 1+*d*_*margin*_/*b*, 1+*d*_*margin*_/*c*), correspondingly. Here *a*, *b* and *c* are the unit cell lengths.
b. In the *extended basic unit cell* we define a grid with a step *h*_*grid*_ (smaller than *d*_*overlap*_) and all grid nodes set to zero.
c. Using the box content table we identify all atoms that belong to this particular box (box *n*) and cycle over them.
d. For each such atom we look up (*t*_*x,n*_, *t*_*y,n*_, *t*_*z,n*_) in the box content table and subtract it from its coordinates; the resulting point belongs to the basic unit cell. For the shifted atom, we define three spheres of increasing radius of 0 < *d*_*overlap*_ < *d*_*contact*_ < *d*_*margin*_ and reassign values of 3, 2 and 1 to all grid nodes inside the respective spherical shell unless the current grid node value is already larger than the new value. This means set number 3 to all grid nodes that have values less than 3 and that are within *d*_*overlap*_ distance to this shifted atom, otherwise we assign 2 to all grid nodes that have values less than 2 and that are closer to this shifted atom than *d*_*contact*_; finally we mark by 1 all zero-valued grid nodes at the distance between *d*_*contact*_ and *d*_*margin*_ (note, mask values can only increase). We emphasize that the grid is defined in fractional coordinates while the distances are calculated in Å; also, grid node values in the mask can only increase.
e. After cycling over all atoms mentioned above, all grid nodes contain integer values, from 0 to 3. Value 3 marks the points with a distance closer than *d*_*overlap*_ to one of the model atoms; an atom from a symmetric copy occurring at this point overlaps with the initial copy. Value 2 marks the points where an atom coming from the symmetric copy will have a contact with at least one atom of the initial copy. Finally, value 1 marks the position of the centers of residues some atoms of which may have a contact; however further checking, atom per atom, is required if this actually happens. Value 0 marks the rest of points where for sure no close atomic contacts may happen.

### 3.2. Checking the symmetry

The steps below are repeated for each symmetry operation *S*_*kl*_ = {*R*_*kl*_; ***T***_*kl*_} defined in §2.2, one by one. For each symmetry operation, we cycle over all residues in the original model as following:

a. For each residue we take its center ***C***_*j*_ calculated as described in §2.4 and apply the symmetry operation

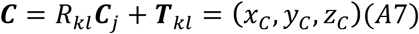
b. Next, we shift the result to the basic unit cell; to do so we define the vector **u** with the coordinates

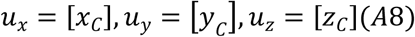

and then calculate

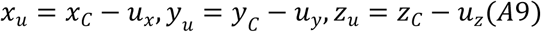
c. We obtain the value of the mask at the grid node closest to (*x*_*u*_, *y*_*u*_, *z*_*u*_). If this value is positive, this residue *j* may have close contacts with the original copy and we analyze the situation in more details as described below:

- For each atom *m* that belongs to residue *j* we recalculate its coordinates as

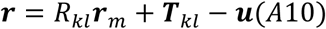
- If the mask at the grid node closest to **r** is equal to 3, this is an abnormal situation highlighting overlapping atoms; unless this atom is on special position (then it shall be processed separately; by default, we simply ignore it) a warning is issued and calculations stop.
- If the mask at the grid node closest to **r** is 2 we accept this residue as the interacting one. We associate the corresponding model transformation as {*R*_*kl*_; ***T***_*kl*_ − ***u*** + ***t***_*n*_. This transformation puts the corresponding symmetric copy close to the original model; that is if we apply this transformation to all atoms of this residue, it will generate a corresponding symmetry copy next to it.
- If the residue is accepted, we do not need to check other atoms of this residue in this position for acceptance. However we need to check, as above, if some of them overlap. If all atoms are checked unsuccessfully, the residue in this position is rejected.

Then we repeat the previous step 3.2c for all 26 model positions around the basic unit cell. For this goal, each of the coordinates of vector **u** is increased or decreased by 1.

As a result, all residues are checked in all symmetry related position around the original model and the contacting residues are identified. Figure A2 summarizes the algorithm.

**Figure A1.**
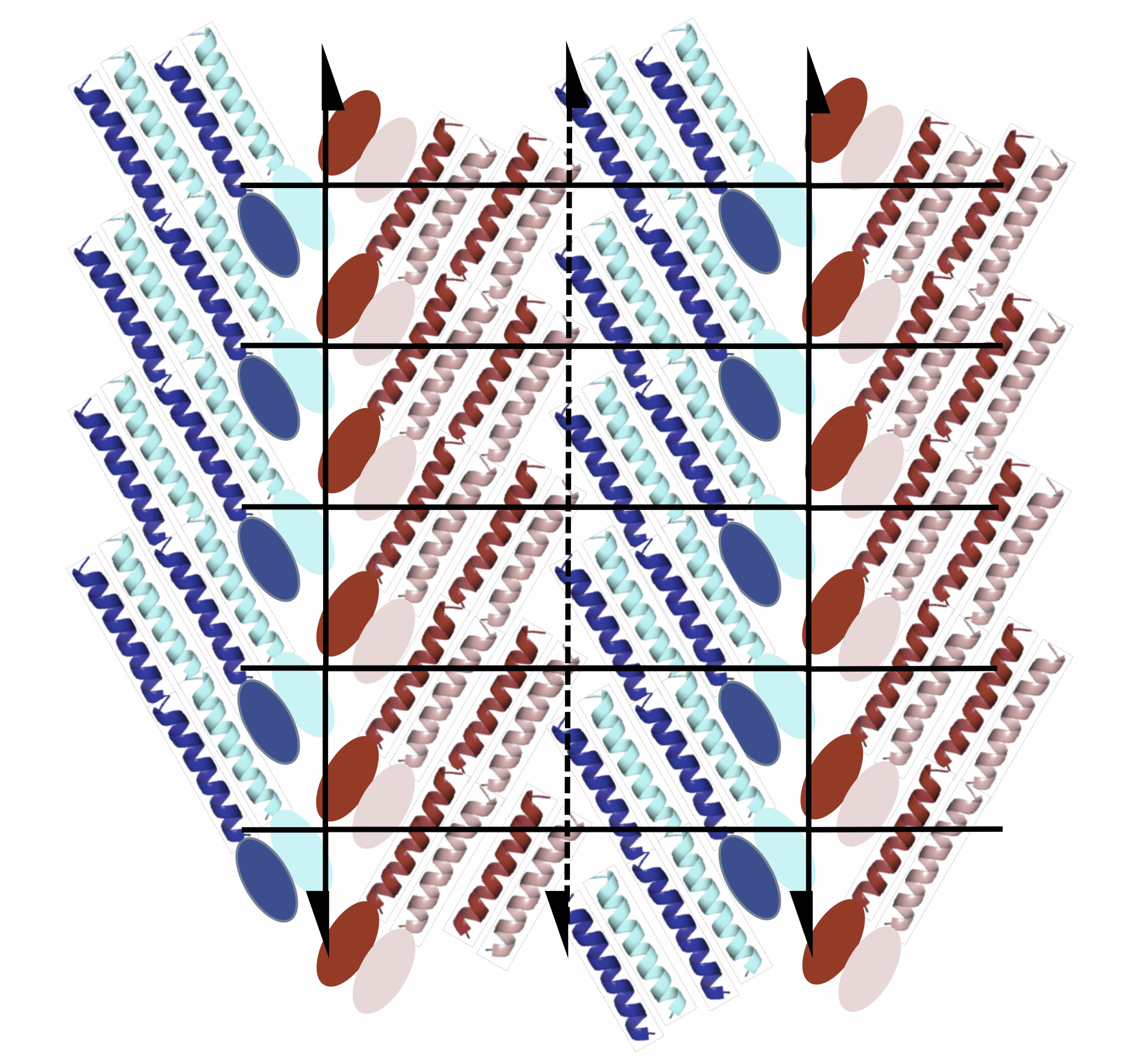
Packing illustration of a crystal with P2_1_ symmetry. Unit cell is a rectangular formed by solid lines (four unit cell shown). A dashed line shows the screw symmetry in the middle of the cell. Each molecule is composed of an elliptic head and a long helical tail marked by the same color. Brown-pink and blue-cyan molecules are related by a local symmetry; brown-cyan and pink-blue are related by a crystallographic one. Each molecule belongs to 4 unit cells, and each unit cell contains the pieces, 4 per molecule, that can be assembled in a whole molecule by translation.

**Figure A2.**
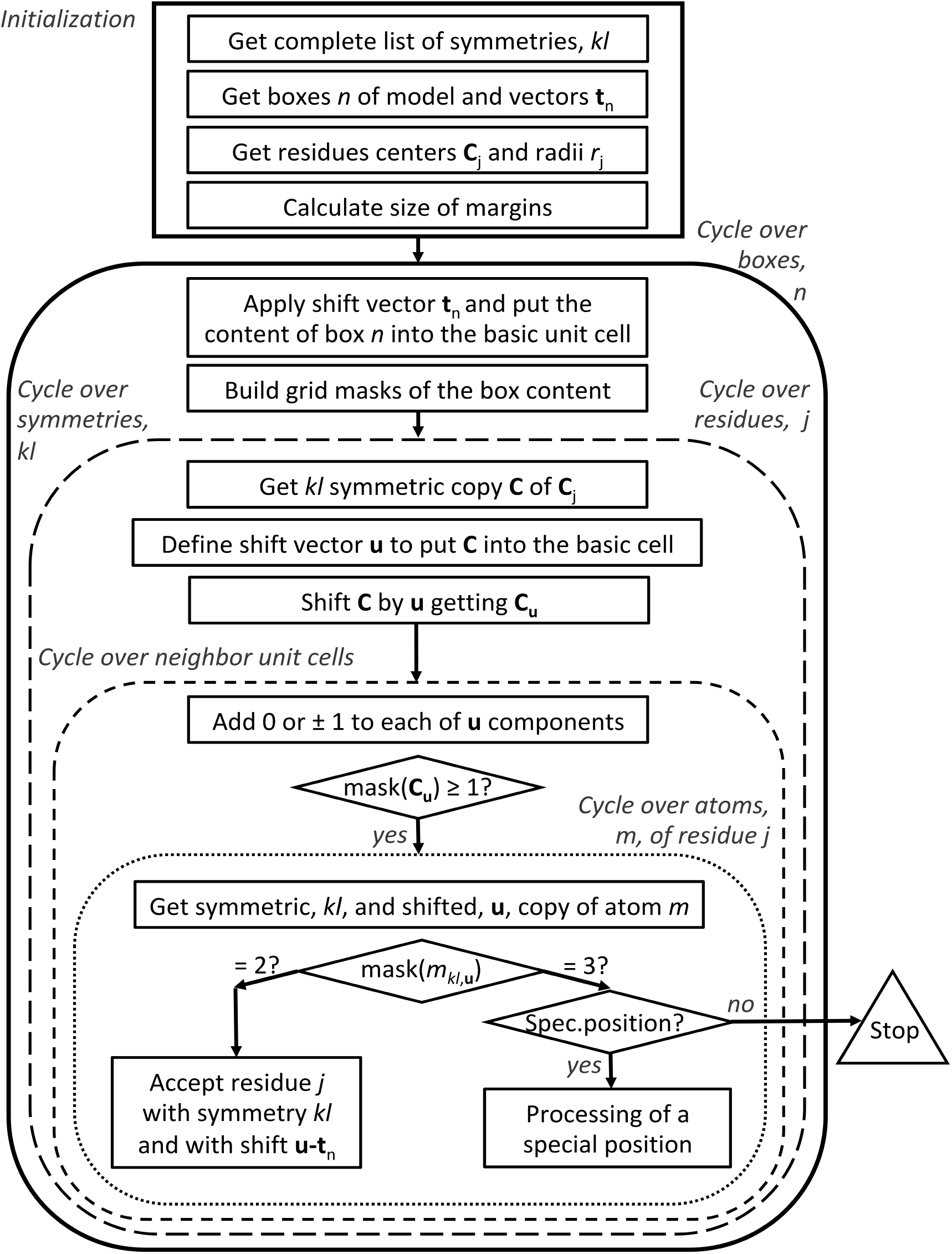
Schematic representation of the process of generating nearest neighbors.

## Footnotes

1 https://www.chemie.uni-bonn.de/pctc/mulliken-center/software/xtb/xtb

2 Based on its definition, the rule-of-thumb interpretation of the Ramachandran plot Z-score (Hooft *et al*., 1997; Beusekom *et al*., 2018) can be summarized as Z<-3 is poor, −3<Z<-2 is suspicious and Z>-2 is good.

